# Bimodal Endocytic Probe for Three-Dimensional Correlative Light and Electron Microscopy

**DOI:** 10.1101/2021.05.18.444466

**Authors:** Job Fermie, Leanne de Jager, Helen Foster, Tineke Veenendaal, Cecilia de Heus, Suzanne van Dijk, Corlinda ten Brink, Viola Oorschot, Lin Yang, Wei Li, Wally Müller, Stuart Howes, Andrew Carter, Friedrich Förster, George Posthuma, Hans Gerritsen, Judith Klumperman, Nalan Liv

## Abstract

Correlative light and electron microscopy (CLEM) can infer molecular, functional and dynamic information to ultrastructure by linking information of different imaging modalities. One of the main challenges, especially in 3D-CLEM, is the accurate registration of fluorescent signals to electron microscopy (EM). Here, we present fluorescent BSA-gold (fBSA-Au), a bimodal endocytic tracer as fiducial marker for 2D and 3D CLEM applications. fBSA-Au consists of colloidal gold (Au) particles stabilized with fluorescent bovine serum albumin (BSA). The conjugate is efficiently endocytosed and distributed throughout the 3D endo-lysosomal network of the cells, and has an excellent visibility both in fluorescence microscopy (FM) and EM. We demonstrate the use of fBSA-Au in several 2D and 3D CLEM applications using Tokuyasu cryosections, resin-embedded material, and cryo-EM. As a fiducial marker, fBSA-Au facilitates rapid registration of regions of interest between FM and EM modalities and enables accurate (50-150 nm) correlation of fluorescence to EM data. Endocytosed fBSA-Au benefits from a homogenous 3D distribution throughout the endosomal system within the cell, and does not obscure any cellular ultrastructure. The broad applicability and visibility in both modalities makes fBSA-Au an excellent endocytic fiducial marker for 2D and 3D (cryo-)CLEM applications.

## Introduction

The ability to characterize the spatial and temporal characteristics of organelles and proteins is crucial in many areas of cell biology. These properties are commonly examined by the use of fluorescence microscopy (FM). FM is highly sensitive, has a large toolbox to simultaneously analyze multiple cellular parameters and can be used to examine the dynamics of processes in live cells. A limitation, however, is its inability to report on the structural context underlying the molecular localization patterns. To obtain this information, electron microscopy (EM) is the method of choice. Its greater resolving power uniquely allows to locate proteins within the cellular context, thanks to its ability to directly visualize membranes and other macromolecular structures. However, specific proteins or structures of interest cannot always be discerned by morphology alone and additional, electron-dense labeling methods are required to visualize them. Various labeling methods are available for EM, such as immunolabeling with colloidal gold^1,2^ or peroxidase-based methods generating osmiophilic precipitates^3–5^. These strategies do have limitations though, as the electron-dense precipitates generated by peroxidase reactions can obscure ultrastructural detail and immunolabels have an inherently limited penetration into specimens^6,7^. The level of penetration can be improved by permeabilization using detergents, but mostly at the cost of structural preservation. A strategy to overcome these drawbacks is correlative light and electron microscopy (CLEM), which integrates the data from FM and EM on a single sample. CLEM uses the large reporter diversity and sensitivity from FM to provide protein localization information, and register this information to high-resolution morphological data, without the limitations of EM labeling.

One of the main challenges in CLEM is to accurately assign the fluorescent label from the FM to the corresponding feature in EM. Retracing specific fluorescent cells or subcellular structures within a large dataset requires reference points that must be easily identifiable in both modalities. These can be naturally formed landmarks such as branching blood vessels and unique cell shapes^8^, artificial marks on the sample support^9–12^, or the sample itself^8,13–15^. Yet, these landmarks are simply too large and do not provide the required registration accuracy to correlate fluorescence signals to individual subcellular structures with nanometer precision. In addition, they may not always be present in quantities sufficient for accurate registration. For these applications, correlation is ideally achieved through artificial fiducials^16–22^. Fiducials are particles easily visible in both FM and EM, which are small enough to not obscure morphological details (<100 nm). A variety of particles have been developed and used in correlative methods, achieving correlation accuracies well below 100 nm ^17–19,23^. These approaches work most efficiently in 2D CLEM applications, where the fiducials are commonly applied to the surface of a substrate, *i.e*. on a coverslip or the formvar layer of an EM grid. However, with the increasing popularity of live-cell and 3D CLEM applications^24^, there is a pressing need for strategies to distribute fiducials *intracellularly* in 3D, which is as of yet unaddressed.

To guarantee accurate registration in 3D CLEM applications, fiducials should be present throughout the entire volume of interest. The endosomal system naturally provides a 3D network of vesicular structures throughout the cell and is easily reached and manipulated from the extracellular environment. This property triggered us to explore the endo-lysosomal system as means to distribute fiducials throughout the cell. Endocytic tracers such as dextran, albumins or nanoparticles conjugated to fluorophores or colloidal gold are commonly used to mark endosomal compartments. One of the most applied endocytic probes, bovine serum albumin (BSA), efficiently labels early endosomes, late endosomes and (auto)lysosomes, and can be conjugated to a variety of fluorophores and colloidal particles^25–27^. BSA conjugates to fluorophores or gold particles have been used in the past for either FM or EM approaches. These conjugates are highly visible and show no cytotoxicity, even after prolonged chase times^27–29^. We reasoned that fluorescently labeled BSA conjugated to electron dense particles could function as an efficient fiducial for 3D CLEM, but no such probe is available to date.

Here, we present an endocytic tracer consisting of 5 or 10 nm colloidal gold stabilized with fluorescently labeled BSA, hereafter named fBSA-Au^5^ and fBSA-Au^10^, respectively. We demonstrate the applicability of fBSA-Au as fiducial marker in a variety of 3D CLEM approaches using Tokuyasu on-section CLEM, resin-embedding 3D-CLEM, and targeted lamella preparation for cryo-EM. We show that the small size of the conjugate enables efficient endocytosis, whereas the high atomic number of the gold colloids ensures good visibility in EM. Importantly, the bimodal nature of the tracer guarantees accurate registration of FM and EM data, by which endocytic compartments containing multiple fluorescent-gold colloids serve as fiducial landmarks. Finally, the uniform size of the gold particles ensures excellent compatibility with immunoEM (double) labeling strategies^30^. These benefits make fBSA-Au a highly useful tool for both 2D and 3D (cryo-)CLEM applications.

## Results

### fBSA-Au particles are stable and monodisperse

As an electron-dense core of the bimodal probe, we chose colloidal gold particles nominally sized at 5 or 10 nm, because of their small size and high visibility in EM. The small size guarantees efficient endocytosis, precise correlations, and compatibility with other immunoEM methods. Using the protocol developed by Slot *et al*.^30,31^, we synthesized monodisperse colloidal particles of different sizes (Figure 1A, D, G). The resulting gold colloids were stabilized with BSA-Alexa555 and purified using centrifugation on a glycerol gradient to remove unbound BSA-Alexa555. This process yielded BSA-conjugated particles of uniform size, with fBSA-Au^5^ averaging 5.8 ± 0.7 nm and fBSA-Au^10^ averaging 8.6 ± 0.5 nm in diameter, as measured in transmission electron microscopy (TEM) (Figure 1A, B, D, E). BSA binding to the particles could not be detected using TEM, likely the result of the low electron contrast of the BSA, which is poorly visible in EM without heavy metal contrasting (Figure 1H).

**Figure 1.**
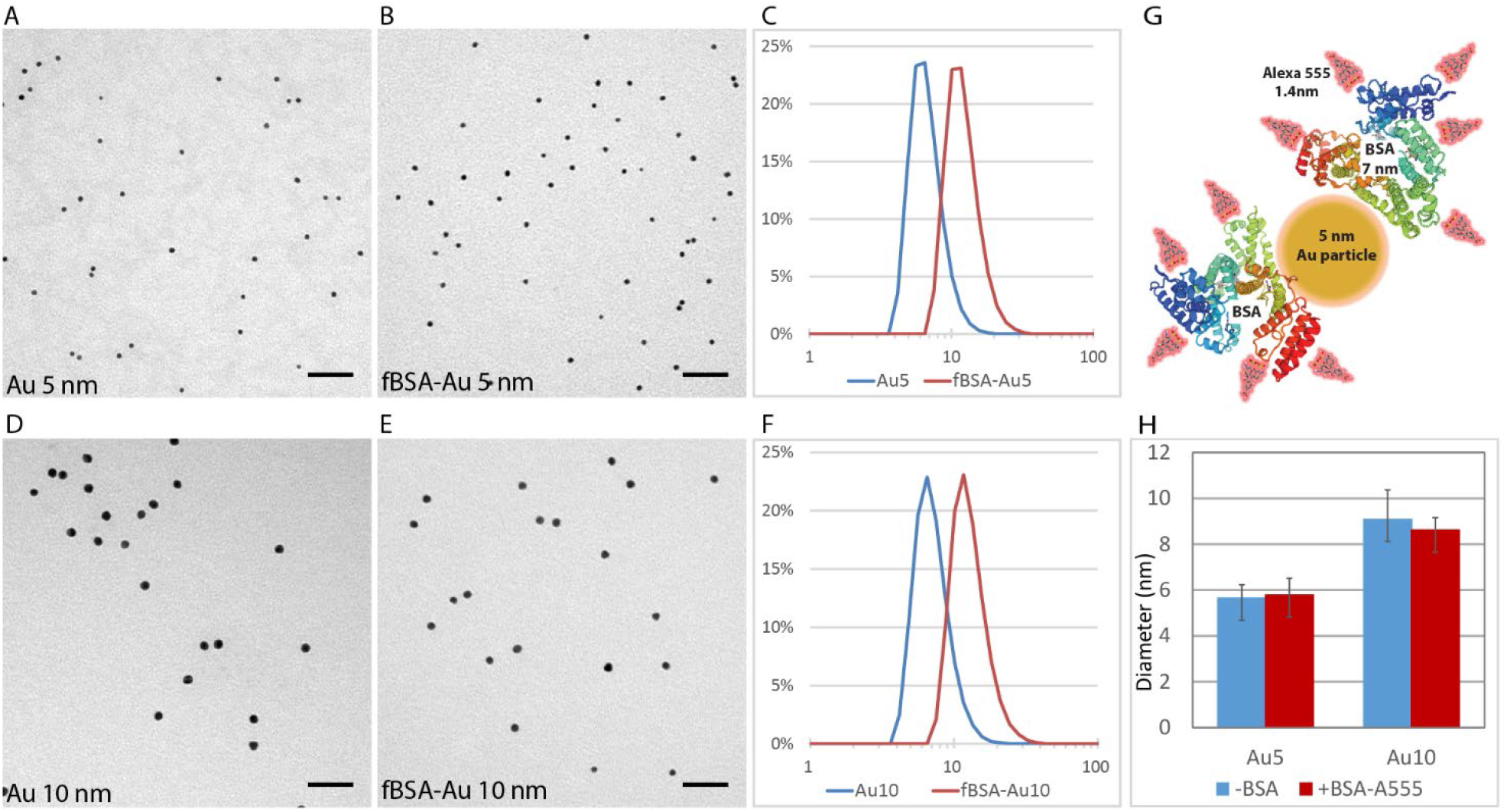
Characterization of synthesized fBSA-Au conjugates. (A, B) Representative TEM micrographs of 5 nm colloidal gold, showing the monodisperse nature of the particles before (A) and after (B) BSA functionalization. (C) DLS measurements showing the size distribution of 5 nm colloidal gold and BSA-Alexa^555^ functionalized particles. Functionalization causes a shift in size distribution, but does not induce larger aggregates. (D, E) Representative TEM micrographs of 10 nm colloidal gold before (D) and after (E) BSA functionalization. (F) DLS measurements showing the size distribution of 10 nm colloidal gold and BSA-Alexa^555^ functionalized particle. Again, BSA functionalization causes a shifted size distribution, but does not induce larger aggregates. (G) Representative schematic showing the relative sizes of BSA (7.1 nm), Alexa555 (1.4 nm), and Au particle (5 nm). (H) Sizing of Au^5^ and Au^10^ particles determined by TEM. Both Au^5^ and Au^10^ are homogeneously sized. Unlike in DLS measurements, the apparent size of the functionalized particles does not increase after BSA functionalization, likely due to the poor visibility of the electron-lucent BSA. Graph depicts mean +/-S.D. *Scale bars: A, B, D, E: 50 nm*.

Stabilization of the gold particles with BSA-Alexa555 resulted in a detectable shift in size distribution, indicating binding of BSA-Alexa555 to the gold particles without forming aggregates of larger sizes. To test clustering behavior, size distributions of the particles were recorded using dynamic light-scattering (DLS) (Figure 1C, F). Here, we found that both the ‘bare’ and ‘functionalized’ particles remain non-clustered in solution. The schematic in Figure 1G shows the relative sizes of a 5 nm Au particle, BSA molecules (7.1-7.5 nm), and Alexa555 molecules with respect to each other^32^. Furthermore, we found that fBSA-Au conjugates remain stable in solution over extended periods of time, showing no signs of clustering or precipitation after several months of storage at 4°C. Together, these data show that colloid gold particles can be functionalized with BSA-Alexa555, after which the resulting conjugates remain monodisperse, uniformly sized and stable in solution.

### fBSA-Au is efficiently endocytosed and transported by the endosomal system

Following characterization of the size and clustering behavior of the conjugates, we assessed the feasibility of fBSA-Au for cellular CLEM experiments. To be useful as an endocytic CLEM probe, the probes should be non-toxic to cells, efficiently endocytosed and brightly fluorescent throughout the experiment.

We first examined the uptake efficiency and localization of internalized fBSA-Au^5^ using FM. After endocytosis, BSA conjugates are predominantly targeted to the degradative path of the endosomal system, where they accumulate in late endosomes and lysosomes^33,34^. We incubated HeLa cells with fBSA-Au^5^ for 3 hours to label the entire endosomal pathway, including lysosomes, the terminal compartments of the endocytic route^27,34^. Following uptake, cells were chemically fixed and immunolabeled for EEA1 and LAMP-1 to mark early endosomal and late endosomal/lysosomal compartments, respectively (Figure 2A). This showed that internalized fBSA-Au^5^ results in the appearance of strongly fluorescent spots that colocalize with both EEA1 and LAMP-1 (Figure 2B). Line profile (Figure 2C) and colocalization analysis (Figure 2D) show that the majority of fBSA-Au^5^ is colocalizing with EEA1 and LAMP-1, only a negligible part (3,8%) is not colocalizing with these early or late endo-lysosomal markers. The data show fBSA-Au^5^ is efficiently taken up by cells and transported through the endo-lysosomal pathway, reaching both early and late endocytic compartments.

**Figure 2.**
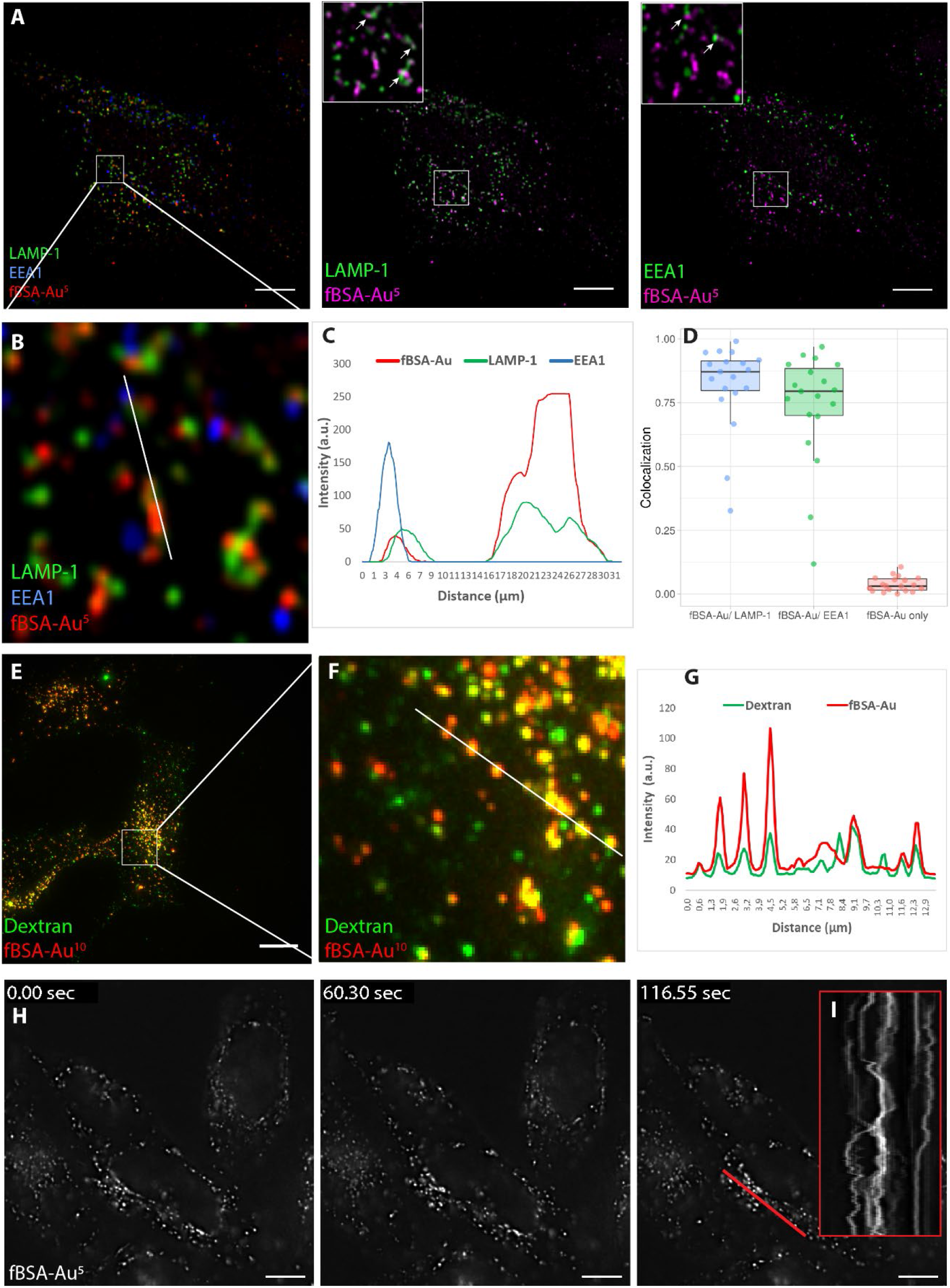
fBSA-Au is a highly fluorescent, efficiently internalized endocytic tracer. (A) Fluorescence images of fixed Hela cells incubated with fBSA-Au^5^ and immunolabeled for EEA1 and LAMP-1. fBSA-Au^5^ is visible in both EEA1 and LAMP1 positive compartments, i.e. throughout the endosomal system. (B) Zoom-in to the white square in (A), (C) depicted line profile, and (D) the colocalization analysis show internalized fBSA-Au^5^ colocalize with EEA1 and LAMP-1. (E-) Fluorescence images of fixed Hela cells incubated with fBSA-Au^10^ and Dextran-Alexa488. (F) Zoom-in and (G) the intensity profile over the depicted lune show fBSA-Au^10^ is readily endocytosed and largely co-localizes with Dextran. (H) Stills from supplementary video 1 of live Hela cells loaded with fBSA-Au^5^. fBSA-Au^5^ has sufficient fluorescence intensity to employ time-lapse experiments with high temporal resolution. (I) Kymograph over the indicated red line shows the x-t scan along the depicted red line. *Scale bars: A-H: 10 μm*.

To examine if fBSA-Au is compatible with the application of commonly used endocytic markers, we incubated HeLa cells with both Dextran488 and fBSA-Au^10^ for 3 hours^35^. Following incubation, cells were fixed and examined by FM. We found that fBSA-Au^10^, like fBSA-Au^5^, is readily taken up resulting in the appearance of brightly fluorescent spots (Figure 2E-F). Line profile shows that the fluorescent signals of dextran and fBSA-Au^10^ majorly colocalized, and on average 80% of fBSA-Au^10^ positive spots also contained Dextran488 (Figure 2G). This indicates that the fBSA-Au probe can be combined with generic endocytic markers and moreover shows that both probes follow the same endocytic route to lysosomes.

Finally, we examined whether the fBSA-Au probes can be applied for live cell imaging, which requires bright and stable fluorescence of the administered probe. After incubating cells for 3 hours with fBSA-Au^5^, coverslips were live imaged with widefield FM. This showed that fBSA-Au^5^ positive fluorescent spots actively moved inside the cells (Figure 2I kymograph along the red line) and remained brightly fluorescent over 250 acquired frames during live-cell imaging (Figure 2H, Supplementary Video 1). We conclude from these combined studies that fBSA-Au is an appropriate probe for FM studies, including live-cell imaging, and which is efficiently taken up by cells and distributes throughout the endosomal pathway.

### fBSA-Au is an efficient fiducial for 2D on-section CLEM

After validating the endocytosis of fBSA-Au^5^ and fBSA-Au^10^, we tested the feasibility of these conjugates for CLEM experiments. First, we tested a 2D on-section CLEM setup using ultrathin Tokuyasu cryosections, the most sensitive method for immuno-EM^9,15,36,37^. Since cryosections in contrast to resin sections show no fluorescent background signal and epitopes are generally well preserved in this approach, they are uniquely suitable for imaging by both FM and EM. CLEM applications using cryosections often involve on-section labeling with a fluorescently tagged antibody that is also marked by colloidal gold^38,39^. The fluorescent labeling is thereby used to select regions of interest (ROIs) for subsequent EM imaging. The fluorescence signal obtained from ultrathin cryosections is limited by the thickness of the sections, which is approximately 70 nm. Therefore, fBSA-Au^5^ fluorescence must be sufficiently intense to be used as fiducial in this approach.

HeLa cells were incubated with fBSA-Au^5^ for 3 hours, after which cells were prepared for cryosectioning and immunolabeling according to our well-established protocol^2,38^. We chose to use fBSA-Au^5^ for these and the following experiments as its small size enables easy distinction from commonly used 10 or 15 nm-sized immunogold labels. Ultrathin cryosections prepared from these HeLa cells were immunolabeled with a primary monoclonal antibody against CD63, a marker for late endosomes and lysosomes, followed by secondary Alexa488-tagged antibody and 10 nm sized protein-A-gold. By FM we found that the Alexa555 fluorophore of fBSA-Au^5^ withstood the preparation steps of cryosections, *i.e*. aldehyde fixation, cryoprotection and plunge freezing, very well. The fBSA-Au^5^ signal was clearly visible in 70 nm thin sections, where it frequently co-localized with CD63 labeling (Figure 3A). We then selected ROIs for imaging in TEM, and registered fBSA-Au^5^ fluorescent spots to EM ultrastructure (Figure 3B, C). We performed an initial correlation based on recognizable features such as nuclei and cell shapes (Figure 3A, B), to find back our cells of interest. Then we used the fBSA-Au^5^ fiducials to zoom in and correlate subcellular structures in FM and EM. We found that Alexa555 labeled compartments in FM always contained 5 nm gold upon EM inspection. The reverse was also valid: gold-containing endosomes and lysosomes identified in EM always contained Alexa555 fluorescence when correlated back to the FM image (see Figure 3D). This showed that fBSA-Au^5^ fluorescence precisely correlates to endosomal compartments containing Au^5^ positive compartments visible by EM.

**Figure 3.**
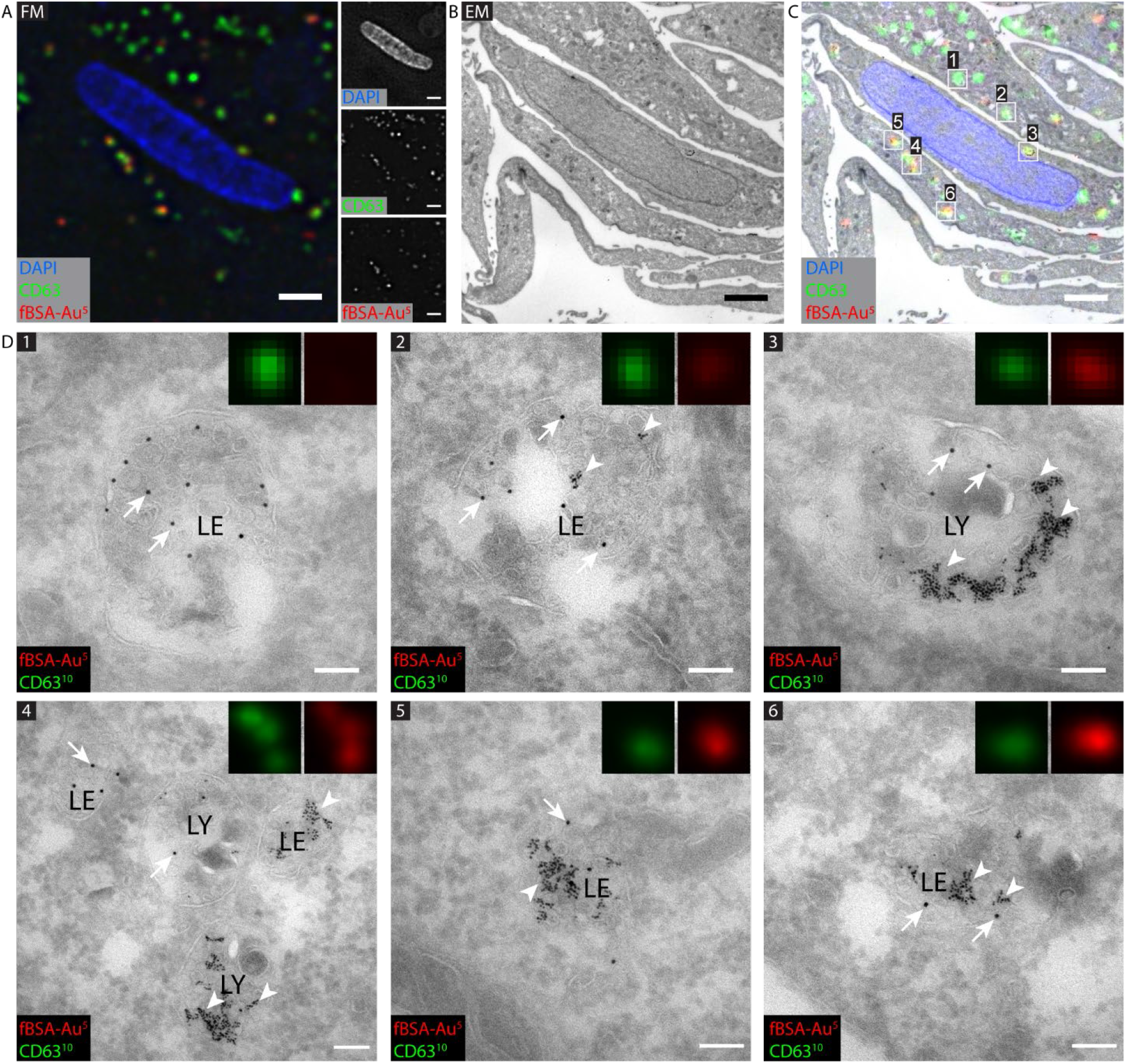
CLEM of CD63 positive, fBSA-Au containing compartments in HeLa cells. (A) FM image of ultrathin cryosection. Region of interest (ROI) with CD63 immuno-labeling (Alexa488 and Au^10^) and Alexa555 fluorescence of internalized fBSA-Au^5^. (B) EM of ROI. (C) Overlay of FM and EM images, showing high registration accuracy between modalities. (D) High magnification EM of fBSA-Au^5^ (arrowheads) containing organelles labeled for CD63 (10 nm gold, arrows). All selected compartments are positive for CD63 (10 nm gold, arrows). Inserts: magnified zoom-ins of CD63 and fBSA-Au^5^ fluorescence of the selected compartments. The intensity of fBSA-Au^5^ Alexa555 fluorescence corresponds to the number of gold particles present per compartment. (1) and (2) Late endosomes (LE) containing many intraluminal vesicles and no (1) or little (2) fBSA-Au^5^ label. (3) lysosome (LY) with clusters of fBSA-Au^5^ gold particles. (4) Several late endosomes and lysosomes containing varying levels of fBSA-Au^5^. (5) and (6) Late endosomes heavily loaded with fBSA-Au^5^ correlating to intense red fluorescence. Scale bars: A-C: 2 μm; D: 100 nm.

Notably, the intensity of the fBSA-Au^5^ fluorescent spots correlated well to the number of gold particles present in a compartment; bright fluorescent spots correlated to compartments with large numbers of gold particles, whereas small, yet readily detectable fluorescent spots correlated to individual particles or small clusters of gold (Figure 3D). This fluorophore-gold correlation is made possible since both fluorescent and gold signal are from the same section, containing precisely the same number of fBSA-Au^5^ particles. To establish the resolution of the correlation approach, we selected the center of fBSA-Au^5^ fluorescence for a given organelle on FM images and the center of the same organelle in EM images^40^. By this we reached sub-organelle accuracy registration. For example, manual picking of 15 pairs of FM and EM fBSA-Au^5^ signals followed by semi-automated correlation with a previously established correlation algorithm, ec-CLEM^40^ (Supplementary Figure 1), resulted in a registration accuracy between 60-130 nm. Importantly, this high accuracy of correlation also enabled us to correlate organelles labeled for CD63 but lacking fBSA-Au^5^ (Figure 3D, panel 1).

These data show that the fBSA-Au^5^ probe correlates with 100% efficiency between the FM and EM modalities, and by both distribution and intensity can be reliably used as a fiducial marker to overlay FM to EM images. Importantly, mapping the constellation of multiple fBSA-Au labeled compartments provides the resolution to correlate fluorescently-labeled structures that lack fBSA-Au, which allows a broad application of the probe to endo-lysosomal as well as other structures. We conclude that its reliable visibility in FM and EM and its compatibility with immunogold labeling makes fBSA-Au^5^ a highly suitable bimodal probe and fiducial marker for 2D section CLEM, with a particularly high accuracy thanks to the inherent wide distribution of particles within the cells.

### fBSA-Au is a suitable fiducial for 3D correlative FM - electron tomography

To extend the applications of fBSA-Au to 3D on-section CLEM applications we next made semi-thin (circa 350 nm) cryosections and prepared these for FM and electron tomography (ET). ET provides a unique tool to view 3D nanometer-scale detail within the cellular context^41^. ET of semi-thin cryosections is a powerful approach for high-resolution 3D ultrastructural analysis of organelles^42,43^.

HeLa cells that were incubated for 3 hours with fBSA-Au^5^ were prepared as stated in the previous section, after which ~350 nm cryosections were prepared. The sections were thawed and immunolabeled for the lysosomal marker LAMP-1 followed by secondary labeling with Alexa488-tagged antibodies and 10 nm sized protein-A-gold. Of note, in contrast to fBSA-Au^5^ added to whole cells for internalization, the antibody-based LAMP-1 labeling is applied onto sections. The antibodies can penetrate into cryosections – as opposed to resin embedded materials – but gold particles mostly remain at the section surface ^6,7^ Hence, there is no direct correlation between the intensities of FM and EM labeling of LAMP-1^44^.

By FM, fBSA-Au^5^ was readily detected as distinct spots that partially colocalized with LAMP-1 (Figure 4A). The overall fluorescence intensity of fBSA-Au^5^ was higher than in 70 nm cryosections (Figure 3 and Figure 4), which corresponds to a higher number of fBSA-Au^5^ particles in the increased Z-volume of 350 nm. We then selected regions with fBSA-Au^5^ fluorescence for correlation to ET. ET image acquisition comprises the tilting of a sample around 1 or 2 axes, after which a 3D reconstruction is generated based on back projection algorithms^41^. Since ET is generally only performed on a small-sized ROI it is of particular importance to select the proper ROI for image acquisition. In semi-thin sections, the increased thickness causes more scattering of electrons and obscures visibility of structures by 2D TEM, which hampers selection of ROIs. The electron-dense gold particles of fBSA-Au^5^ were very well visible already by 2D TEM, which greatly facilitating the retrieval of an ROI. After identifying the proper ROIs, tilt-images were collected by ET (Figure 4B-D), in which individual organelles were registered from FM to 3D EM. The correlation accuracy of this approach ranged between 60 and 200 nm as determined using ec-CLEM, which is at the same level as for 2D CLEM. These data show that in semi-thin sections the fBSA-Au^5^ probe can be used both at the mesoscale – to find back cells and ROIs – and at the nanometer level, to correlate individual organelles.

**Figure 4.**
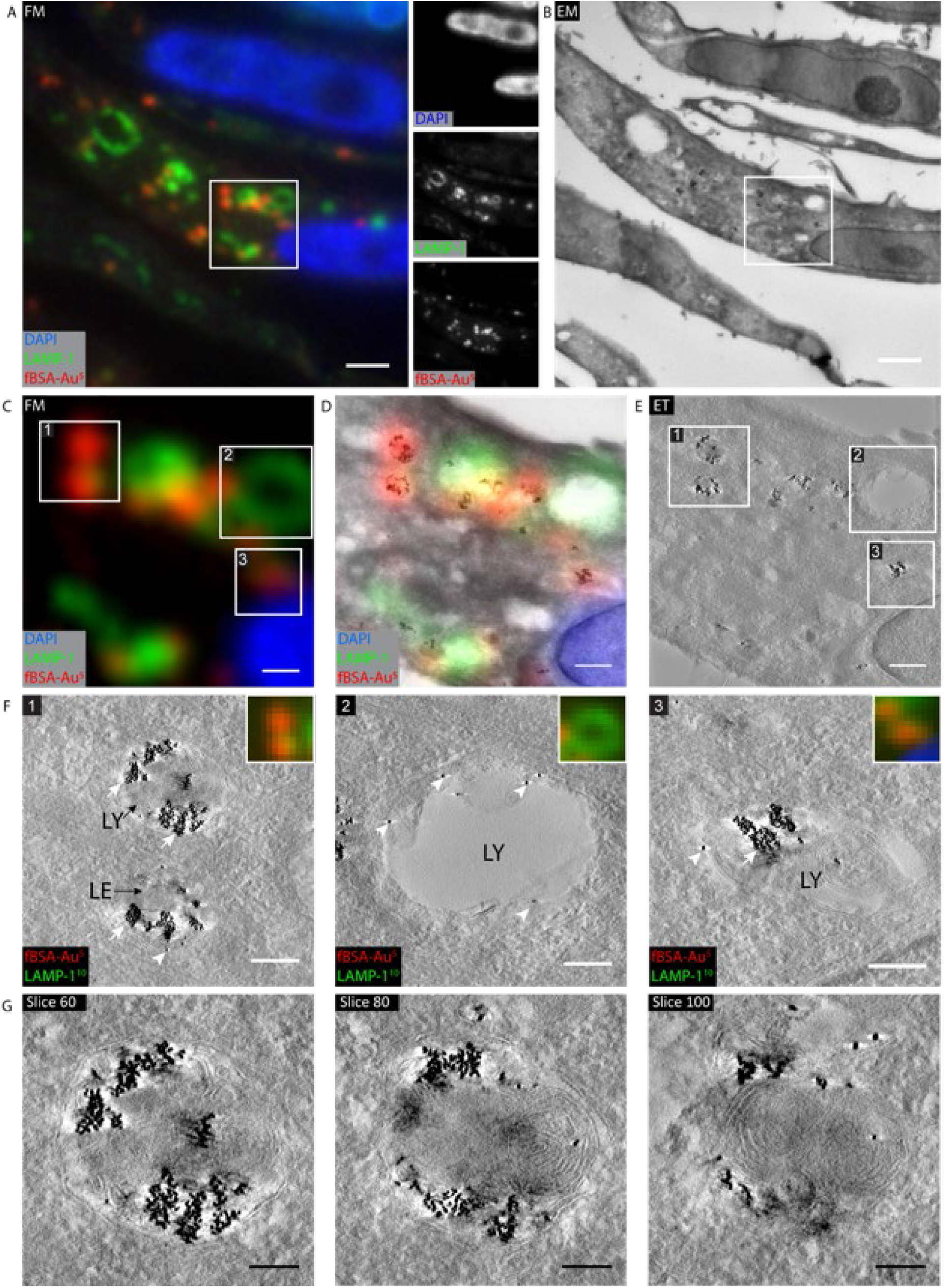
3D CLEM of 350 nm cryosections using electron tomography. Hela cells were incubated with fBSA-Au^5^ fiducials (3 hours) and immunogold labeled for LAMP-1 (10 nm gold). (A) FM image of a 350 nm thick cryosection with the selected ROI for ET highlighted by the white box. Insets show separate channels for the used fluorophores. (B) EM of same region as shown in (A). At this magnification, clusters of endocytosed fBSA-Au^5^ gold particles are visible, allowing rapid correlation from FM to EM. The ROI selected for ET is shown in the white box. (C) Magnified crop from (A) showing the ROI selected for ET. Numbers refer to the same spots as shown in E and F. (D) Zoom-in to ROI with the fluorescence information overlaid. fBSA-Au^5^ fluorescence strictly corresponds to fBSA-Au^5^ gold particles. (E) Virtual slice from tomogram highlighting selected organelles. (F) Zoom-ins of selected organelles. Panel 1: late endosome (LE) and lysosome (LY) with large amounts of fBSA-Au^5^ gold particles (white arrows) and minimal LAMP-1 labeling (white arrowheads). Panel 2: LAMP-1 labeled lysosome devoid of endocytosed fBSA-Au^5^. This slice is from the surface of the section, showing LAMP-1 representing gold particles that do not penetrate into the section. Panel 3: LAMP-1 labeled lysosome abundantly filled with endocytosed fBSA-Au^5^ gold. (G) Virtual sections through the lysosome shown in panel 1 of (F), showing distribution of gold throughout compartment *Scale bars: A,B; 2 μm; C-E: 500 nm; F: 200 nm; G: 100 nm*.

After tomogram reconstruction, we examined the morphology of the correlated structures. ROI 1 (Figure 4C, D, E, F) showed a collection of fBSA-Au^5^ particles present in a late endosome and a lysosome. Multiple virtual slices through the lysosome showed that fBSA-Au^5^ is evenly distributed throughout the organelle (Figure 4G). ROI 2 contained a large, LAMP-1-positive organelle negative for fBSA-Au^5^. The corresponding ET image shows only sparse LAMP-1 gold-labeling on the section surface. Thus, despite the absence of fBSA-Au^5^ and sparse immunogold labeling, these organelles could be marked as LAMP-1 positive as detected by FM labeling registered to EM. These data imply that the intensity of the FM signal and the accuracy of the correlation procedure overcomes the need to label fluorescently tagged antibodies with an additional gold tag. ROI 3 showed a typical lysosome containing ample fBSA-Au^5^ particles. Note that the 5 nm gold particles are visible throughout the volume of the tomograms (Figure 4F, G), thanks to the internalization of fBSA-Au^5^ prior to fixation. The intracellular, 3D distribution of fBSA-Au^5^ allowed correlation of structures throughout the section (Figure 4G), which is an important improvement over fiducials that reside on the surface of sections.

Together, these findings show that endocytosed fBSA-Au^5^ is a powerful fiducial marker for 3D CLEM, showing high visibility and enabling fast retracing of ROIs with high registration accuracy. Moreover, the sensitivity of fluorescent labeling combined with the high correlation efficiency overcomes the need for gold labeling of antibodies.

### fBSA-Au as an endocytic fiducial for correlative live-FM and electron tomography of resin-embedded cells

Many CLEM approaches use resin embedding for EM. In these applications, the flow of experiments generally involves recording of fluorescence signals in live or fixed 3D samples, i.e. not sections as above, after which the sample is infiltrated/stained with heavy metals for increased EM contrast and embedded in epoxy or acrylic resin. Then (serial) ultrathin or semi-thin sections are made, which are collected for (serial) TEM or ET and correlated to the FM data. This type of correlative approach is sensitive to distortions between FM and EM datasets, since staining, dehydration and embedding steps are performed after FM imaging. 3D CLEM approaches (serial sectioning and/or ET) introduce an additional level of challenge in the z-dimension, due to the disparity in axial resolution between FM and EM. We reasoned that fBSA-Au fiducials due to their unique 3D distribution would significantly ease such 3D CLEM correlations and tested this by making ETs of serial semi-thick sections.

HeLa cells grown on patterned glass coverslips were incubated for 3 hours with fBSA-Au^5^, fixed and imaged in the FM. Cells were recorded relative to the pattern on the coverslips to facilitate successive X-Y correlation to the EM, a procedure introduced by Polischuk *et al*.^11^ and commonly used in volume-CLEM applications ^45,46^. The volume-CLEM approach is outlined in Figure 5A. In short, a fluorescent Z-stack of the cell with 200 nm intervals was collected,then the sample was contrasted with heavy metals, embedded in epoxy resin, after which 250 nm serial sections were collected for EM. Using the fluorescent Z-stack, the 250 nm EM section bearing the organelles of interest was estimated, imaged, and correlated to the FM data. Figure 5 shows a differential interference contrast (DIC) image from a selected cell (Figure 5B), overlayed with the fluorescence max intensity projection from endocytosed fBSA-Au^5^ (Figure 5B-C) and a low magnification overview of the corresponding 250 nm EM section (Figure 5D). We selected perinuclear regions for FM-EM correlation, since these are relatively thick and best demonstrate the 3D cellular distribution of endocytosed fBSA-Au^5^. We used the information from the FM Z-stack to assess the distance between an ROI and the bottom of the coverslip. This guided us to select the correct section to acquire high-resolution tomograms of the fBSA-Au^5^ containing compartments (method outlined in Figure 5A).

**Figure 5.**
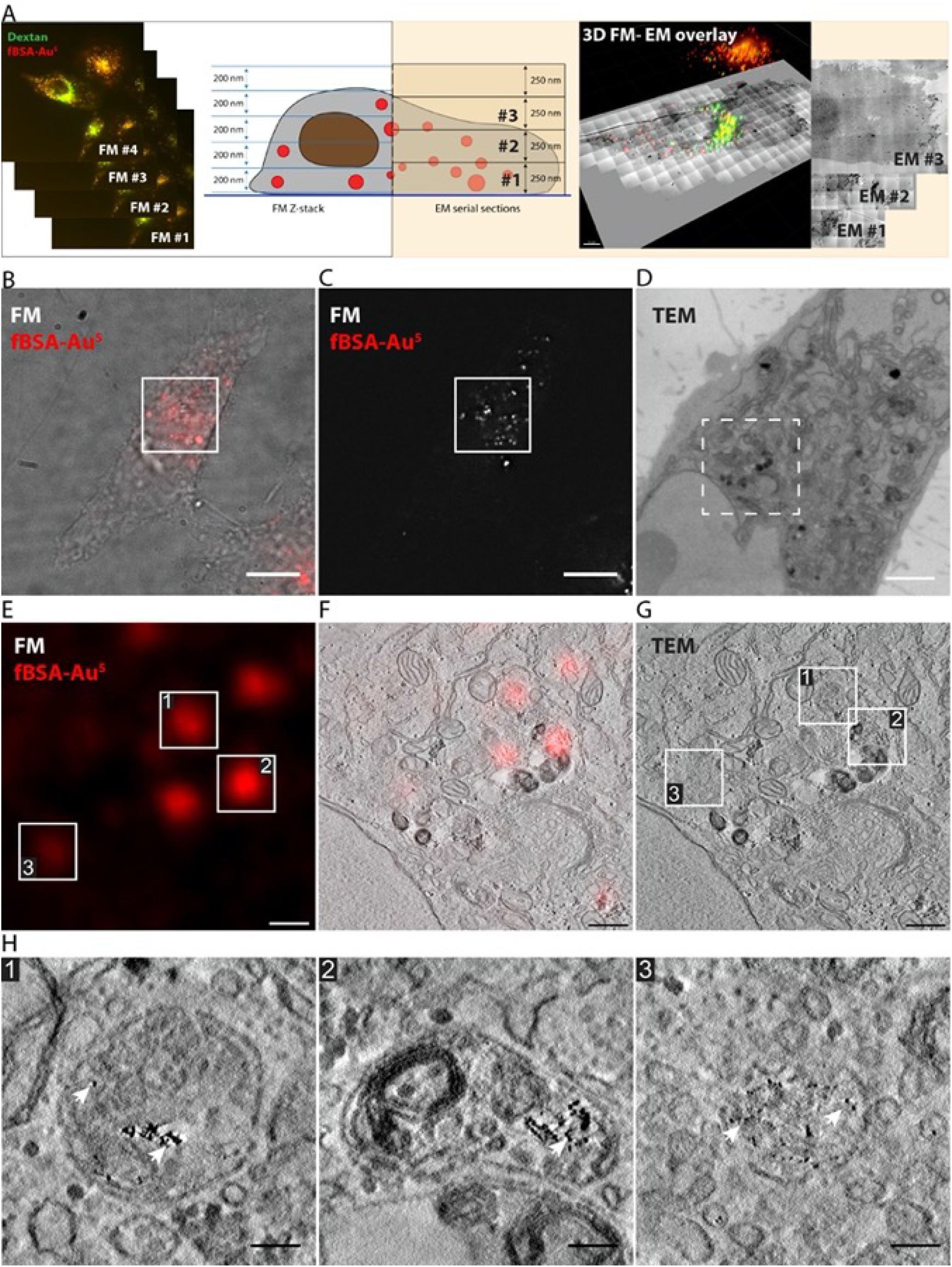
fBSA-Au serves as a bimodal endocytic probe for CLEM using pre-embedding fluorescence and resin sections. HeLa cells were incubated for 3 hours with fBSA-Au^5^. (A) Schematic of the imaging strategy employed. A fluorescent Z-stack with 200 nm intervals is collected after fixation but prior to resin embedding. After resin embedding, 250 nm thick sections were cut for ET. The depth (z-plane) bearing the organelles of interest was estimated based on the fluorescent Z-stack, and the corresponding section was imaged in ET. (B) DIC image of a cell with fluorescence of fBSA-Au^5^ overlayed in red. (C) Fluorescence signal of fBSA-Au^5^ shown in A. ROI for CLEM is highlighted with a white box. (D) TEM micrograph of a 250 nm thick section showing the ROI from B and C. The ROI for ET is indicated in the dashed white box. (E) Fluorescence signal corresponding to the ET ROI with numbered spots of interest. (F) Virtual slice from tomogram overlaid with fluorescence data. (G) Virtual slice from the tomogram showing the 3 selected organelles. (H) Magnified virtual slices of the selected organelles, containing fBSA-Au^5^, visible by the 5 nm gold particles (arrows). Organelles 1 and 2 are late endo-lysosomes, while organelle 3 is a late endosome. Scalebars: B, C: 10 μm; D: 2μm; E-G: 500 nm; H: 100 nm.

In the tomograms, we could easily distinguish the individual organelles selected in FM (Figure 5E, F). Most fluorescent spots correlated by ET were endo-lysosomal compartments containing clusters of gold particles (Figure 5G, H). We also found faint fluorescent spots that correlated to endosomal organelles containing only few gold particles (Figure 5H, organelle 3). This highlights the strong signal of the fBSA-Au^5^ probe. We conclude that endocytosed fBSA-Au is highly suitable for resin-based 3D CLEM approaches: first by providing the information required to select the correct Z-layer for ET imaging, and second by using fBSA-Au as fiducial marker to provide high correlation accuracy in 3D.

### fBSA-Au as a fiducial marker to mark endo-lysosomes for cryo-electron tomography

Cellular cryo-ET is an emerging technique for determining the 3D structures of molecules with sub-nanometer resolution^47^. Vitrification of cells keeps molecules in their near native state and cryo-ET provides high resolution *in-situ* images of molecules within the context of the cell. However, the lack of efficient labels for cryo-ET and the low level of contrast makes it challenging to select subcellular ROIs for cryo-ET image acquisition^48 49^. Cryo-FM screening prior to cryo-ET is a promising way for identification of ROIs, which can then be re-located and targeted for imaging by cryo-ET^50^. We postulated that fBSA-Au would be a highly suitable tool to mark endo-lysosomal organelles by cryo-FM and serve as tool to pre-identify and select regions to collect tomograms by cryo-ET.

For these experiments we used cultured, differentiated neurons (dorsal root ganglion (DRG) neurons), since the <500nm thin axonal projections (transparent for the electron beam) allow for direct cryo-ET without the need to prepare lamella^51^ (Figure 6). We incubated DRGs for 2 hours with fBSA-Au^5^ and then performed the required washing and vitrification steps (see Methods for detailed procedure). By cryo-FM, the red fluorescence of fBSA-Au^5^ was visible in athin region of the axon as a collection of small fBSA-Au^5^ puncta (Figure 6A&B, yellow rectangle inset and arrows). There were also some bright fluorescent patches on the empty grid surface (white arrowheads in Figure 6A), which likely correspond to clusters of fBSA-Au^5^ adhering to the laminin coat required for neuronal growth, since no such patches were observed on glass or uncoated or fibronectin coated grids. We then used the presence of axonal fBSA-Au^5^ labeling to navigate to the axonal regions in the TEM and collect cryo-ET data. The reconstructed tomograms collected of similar ROIs showed various endo-lysosomal compartments of which approximately half contained fBSA-Au^5^. The high contrast of the fBSA-Au particles allowed their unambiguous identification within the cryo-ETs. The gold appeared as single particles in small vesicles (Figure 6C, see also Supplementary Video 2: Tomogram of Figure 6C), tubules and early endosomes (Figure 6D, E), and was more clustered in later endo-lysosomal organelles (Figure 6F, see also Supplementary Video 3: Tomogram of Figure 6F) and (Figure 6G, see also Supplementary Video 4: Tomogram of Figure 6G). These data show that endocytosed fBSA-Au is visible in both cryo-FM and cryo-ET and can be applied to mark ROIs for imaging by cryo-ET.

**Figure 6.**
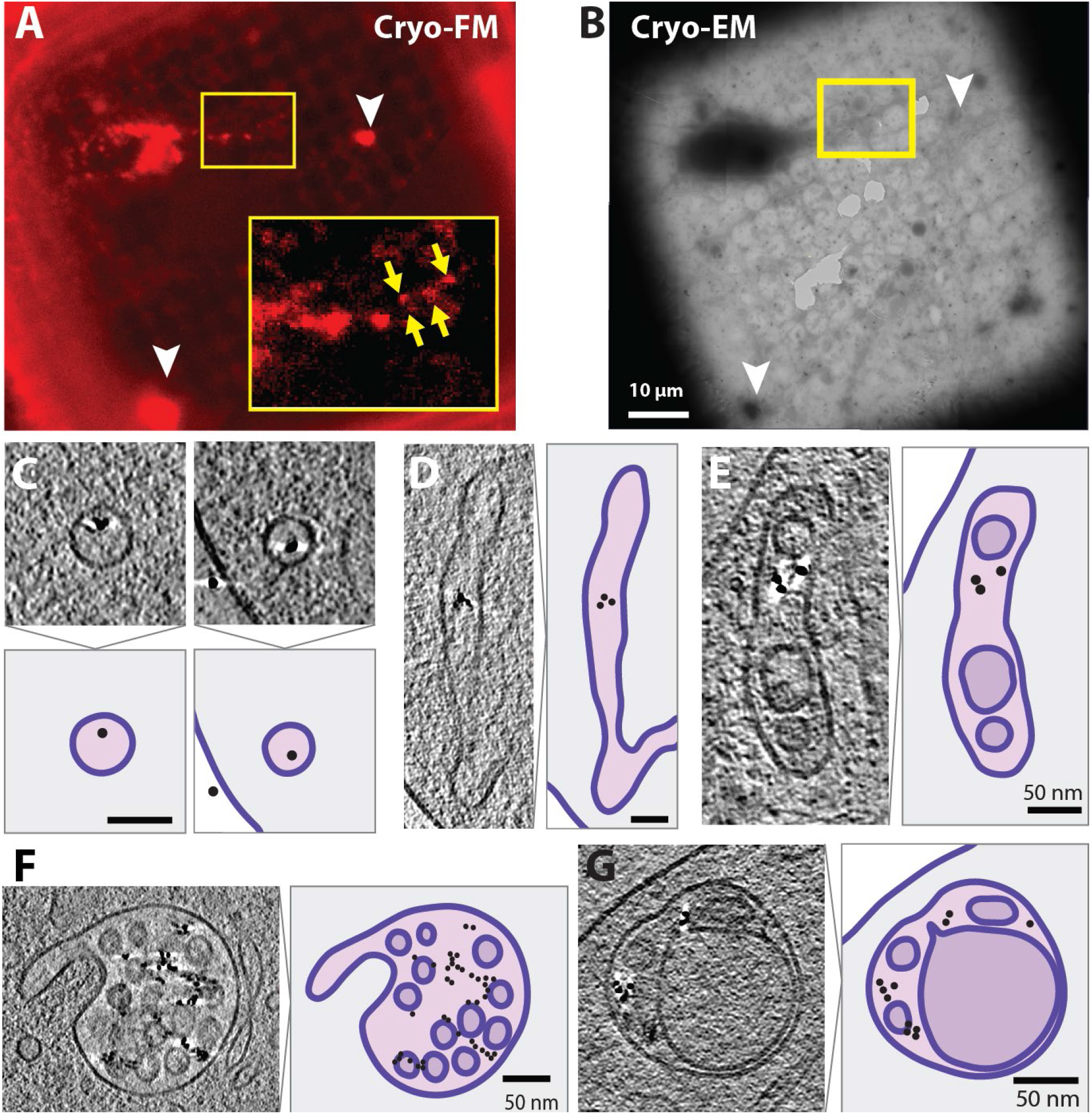
fBSA-Au as a bimodal endocytic probe for cryo-ET. (A) Cryo-FM and (B) cryo-EM images of DRG neurons grown on EM grids. Red signal in cryo-FM originates from the Alexa555 groups of fBSA-Au^5^. Yellow arrows point to less intense fluorescent spots corresponding to small endo-lysosomal organelles in the thin parts of axons. White arrowheads point to dense clusters of fBSA-Au^5^ adhering to the laminin coated grid. (C) Small vesicles in cryo-ETs of DRG neurons containing fBSA-Au^5^ gold particles. Model for each example is shown below, with lipid bilayer shown in purple, vesicle lumen in light pink and fBSA-Au^5^ shown as black circle. Neuron cytoplasm is shown in grey, region outside of the cell is white. (D) As in C, for tubular structures (E) As in CD, for an early endosome. Lumen of internal vesicle is shown in light purple. (F) As in CE, for a multi-vesicular body. (G) As in FC, for a late endosome/lysosome. Scalebars: A,B: 10 μm; C-G: 50 μm.

### fBSA-Au as a fiducial marker for lamella preparation for cryo-ET

Unlike axons, most biological samples are too thick to perform direct cryo-ET. Instead, 100-200 nm lamellae need to be prepared first to enable cryo-ET imaging^52^. In short, for this procedure cryo-preserved cells are milled in a cryo-FIB-/SEM to create lamellae suitable for cryo-ET imaging. Cryo-FIB milling is the most effective thinning method to date, but imaging of specific biomolecular processes requires their localization of ROIs that measure ~1μm laterally and 200-300nm in z in larger cells. Especially for rare cellular events or structures there is no routine method assuring that the structure of interest is present in the prepared lamella. Cryo-FM is used to identify an ROI for cryo-ET. In general, fluorescently labeled cells (e.g. expressing a GFP-tagged protein) are cultured on TEM grids, vitrified, imaged by cryo-FM and transferred to a cryo-FIB-SEM, where the position for lamellae milling is selected in the x-y plane (laterally) based on the cryo-FM data, and the z-position (axially) estimated to the best accuracy allowed by the FM data. Once processed, the lamellae are transferred to a cryo-TEM for imaging. This approach maximizes targeting of the intended ROI and reduces imaging of areas that do not contain structures of interest^48,53^, increasing the success rate of the method while reducing beamtime. Fiducial markers with a homogeneous 3D distribution throughout the intracellular volume are a highly powerful tool to mark ROIs for cryo-ET. Hence, as final application case, we here test the performance of fBSA-Au in a cryo-ET CLEM workflow, using the fBSA-Au fiducial particles to target lamellae preparation and identify targets within the prepared lamellae.

Human bone osteosarcoma epithelial (U2OS) cells were cultured on Dynabead (1 μm sized traditionally used fiducials) treated TEM grids, incubated with fBSA-Au^5^ particles for 3 hours and vitrified. A fluorescent z-stack showing the position of the Dynabeads (green channel) and fBSA-Au^5^ particles (red channel) was collected in a spinning-disk FM equipped with a cryo-module ([see Supplementary Video 5 for the z-stack of the cell shown in Figure 7](https://surfdrive.surf.nl/files/index.php/s/wpiZM1YFQXOuCrX)). A maximum intensity projection of the fBSA-Au^5^ particles present in the z-stack is shown in Figure 7A. After 3D cryo-FM imaging, the grids were loaded in the cryo-FIB-SEM. First, the FM and SEM overview images were correlated by the distribution of the Dynabeads, providing the x-y-z alignment prior to milling (Figure 7B). Then, using the fluorescent z-stack from the fBSA-Au^5^ particles (Figure 7C), a lamella (z) position was selected and prepared as described^54^. The SEM image of the prepared lamella is shown in Figure 7D, overlaid with the fluorescence of the fBSA-Au^5^ particles. The prepared lamella was then transferred and imaged in a 200 kV cryoEM. The lamella overview images from cryo-EM were correlated to the cryo-FM data showing the localization of the fBSA-Au^5^ particles (overlay shown in Figure 7E). We then recorded high magnification images (Figure 7G and 7H) of the correlated organelles (indicated with green and blue squares in Figure 7E and 7F) showing endocytosed fBSA-Au^5^ particles in late endosomes and lysosomes. Notably, we managed to follow a single fluorescent compartment (depicted with a red arrow) throughout the complete cryo-CLEM procedure, and recorded tomogram ([Supplementary Video 6: Tomogram Figure7](https://surfdrive.surf.nl/files/index.php/s/ngRlcOEFiMkypbv), corresponding to the red arrow in Figure 7).

**Figure 7.**
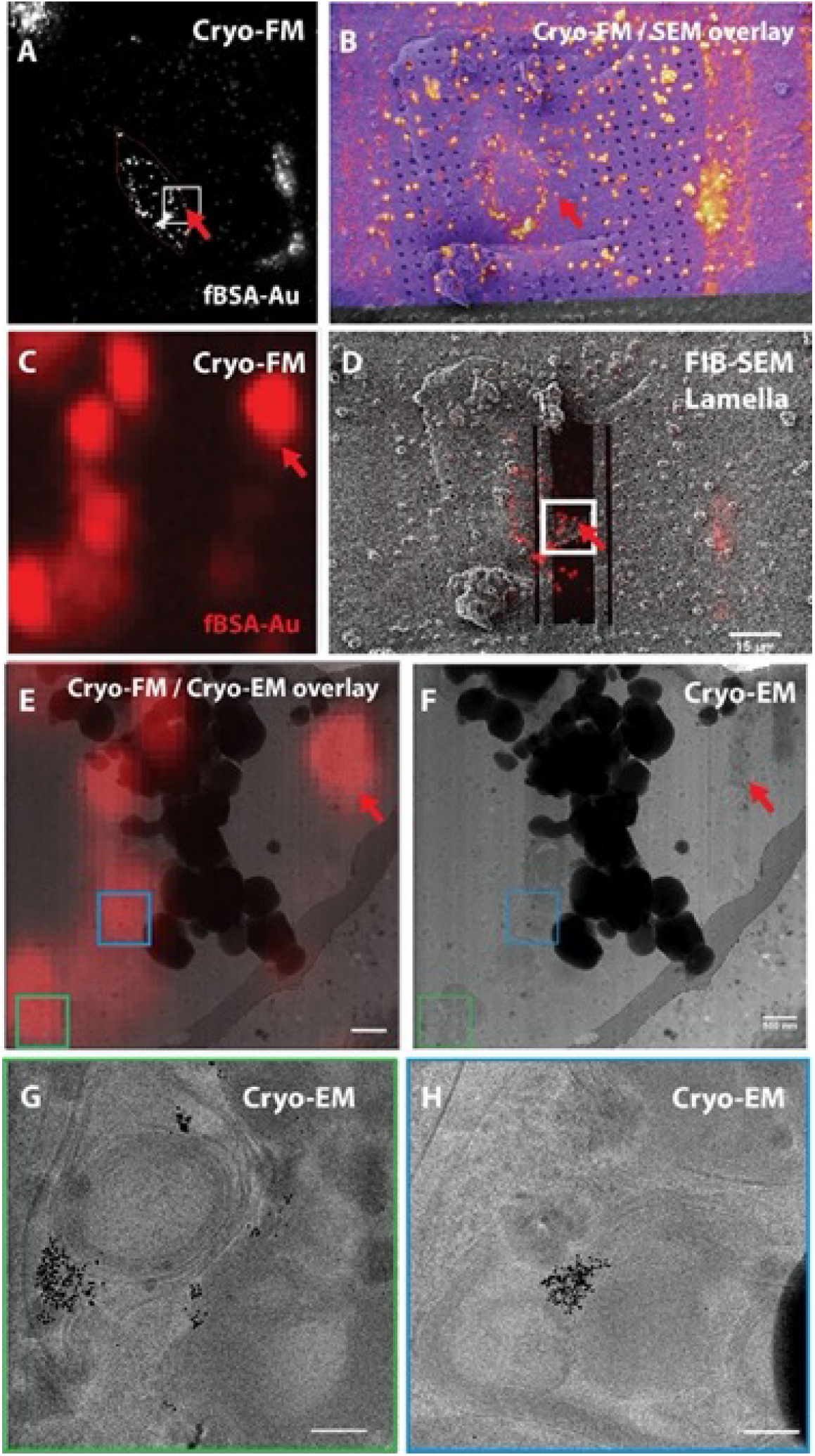
fBSA-Au as a fiducial marker for cryo-CLEM. (A) Red channel maximum intensity projection of the cryo-FM z-stack of the selected U2OS cell. The red signal in cryo-FM originates from the Alexa555 groups of fBSA-Au^5^. (B) Green channel cryo-FM image (Dynabeads) overlayed to the SEM overview. (C) Zoom-in cryo-FM image of the ROI showing the organelles bearing fBSA-Au^5^ selected for lamella preparation. The ROI is also depicted with a white square in (A). (D) Post milling SEM image of the prepared lamella overlayed with the corresponding individual z-stack image of the FM data. (E) Overlay of cryo-FM signal of fBSA-Au^5^ present in the corresponding organelles in the lamella and cryo-TEM images. (F) Overview cryo-TEM image of the lamella. Organelles imaged with higher resolution are depicted with green and blue squares; and shown in (G) and (H) respectively. (G) A lysosome bearing fBSA-Au^5^ in its lumen. (H) Another lysosome bearing fBSA-Au^5^. *Scale bars: B,D: 15 μm; E,F: 500 nm; G,H: 100 nm*.

These data show that endocytosed fBSA-Au can be used as 3D fiducial marker in a cryo-CLEM workflow. The 3D distribution of the particles within the endo-lysosomal system makes them very suitable to select regions for lamella preparation based on the cryo-FM signal. The strong contrast of the particles in cryo-TEM makes their visualization easy, and increases the accuracy in tomogram reconstruction to visualize fine structural elements. For example, reconstructions of fBSA-Au^5^ endosomes readily show contact sites between endosomes and cytoskeleton ([Supplementary Video 7: Tomogram of a microtubule-late-endosome connection](https://surfdrive.surf.nl/files/index.php/s/fmlT9i7ZgPvrZOp)).

## Discussion

Bimodal fiducial markers are of great importance for correlative microscopy methods since they enable an accurate registration of fluorescence information over EM morphology ^16,17,55^. Here, we introduce fBSA-Au, a conjugate of AlexaFluor-labeled BSA and 5 or 10 nm sized colloidal gold particles, and describe its application as a bimodal endocytic probe and 3D fiducial marker. Endocytic markers are an attractive solution for correlative fiducials, since they efficiently distribute throughout cells, resulting in a well-defined 3D pattern of landmarks usable for correlation. We demonstrate that fBSA-Au^5^ and fBSA-Au^10^ are stable in solution, efficiently endocytosed, non-toxic, brightly fluorescent and with good electron contrast for detection in TEM. Furthermore, the probe is versatile, since it can be synthesized with differently sized gold particles and distinct fluorophores. We have successfully generated fBSA-Au^5^ and fBSA-Au^10^ particles with Alexa488, Alexa555, Alexa 647, Texas Red and Tetramethylrhodamine (TRITC) fluorophores. We show that endocytosed fBSA-Au is taken up effectively by cells, colocalizes with markers of early and late endosomal compartments (EEA1, CD63 and LAMP1), and reaches the same population of organelles as do established endocytic tracers such as Dextran or fluorescent BSA.

The properties of fBSA-Au render it broadly applicable for 2D and 3D CLEM, and compatible both with resin EM, cryosectioning, and cryo-EM approaches. We show that fBSA-Au provides ample fluorescence signal for imaging in intact cells in both live and fixed conditions, thanks to the buildup of multiple fBSA-Au particles into the confined space of endosomes and lysosomes. In addition, fBSA-Au is readily detectable even in FM of ultrathin (70 nm) Tokuyasu cryosections. The high electron contrast and uniformly sized gold particles make them easily visible in TEM and reliable reference points to correlate data from FM and EM, in both 2D and 3D. Moreover, since the gold particles in the fBSA-Au’s are of well-defined size, their use is compatible with immunogold labeling, which is especially valuable when using cryosection CLEM. We have demonstrated the use of fBSA-Au in TEM based CLEM, but we envision that fBSA-Au is also compatible with SEM-based CLEM approaches, using 3D EM techniques like FIB-SEM, serial blockface scanning EM (SBF-SEM) and array tomography. SEM-based 3D methods enable the examination of larger volumes than ET, but with less resolution^56^, which likely necessitates the use of fBSA-Au^10^. The excellent visibility in EM ensures compatibility with current and future automated CLEM procedures and registration software ^40^-^57^-^58^. Combining the uncompromised FM and EM properties, we found that the probe functions well for CLEM, enabling live-cell imaging and easy registration of data in both 2D and 3D applications.

The examples provided here show CLEM of endo-lysosomal organelles, but the bimodal visibility of fBSA-Au is also highly usable for CLEM studies of other structures. Endosomes containing fBSA-Au can be used as 3D reference points with a unique spatial distribution to register FM data from other structures of interest beyond the endo-lysosomal system to their corresponding ultrastructure with a correlation precision comparable to organelles bearing fBSA-Au particles (Figure 3, 4). Since nearly all cell types show a significant level of endocytosis, with the exception of erythrocytes^59^, the use of endocytosed fBSA-Au is widely applicable to a large variety of cellular and tissue models. The applicability also extends to live-cell CLEM approaches, since the endocytosed fiducials are visible during every step of correlative imaging, all the way from live cells to the final tomograms (Figures 2,5). Bimodal nanoparticles are a useful tool in cell biology, either as bimodal fiducial for CLEM or as an endocytic probe to mark endo-lysosomal organelles.

Previously, we and others introduced fluorescently labeled silica nanoparticles as bimodal endocytic tracers and fiducials for both FM and EM^16,27^. These particles provide easy correlation, and perform well as extracellular fiducials for CLEM. However, their relatively large size limits the efficiency of endocytosis, and may obscure morphological features within endosomes, hampering their ultrastructural identification. The fBSA-Au probes presented here are significantly smaller (5-15 nm diameter vs 60-100 nm for silica particles) which allows higher levels of endocytosis while the visibility of morphological features in endosomes is retained. Quantum dots^23,60,61^ and nanodiamonds^62,63^ are another type of bimodal nanoparticles used in cell biology and CLEM approaches. Like fBSA-Au, they can be functionalized with physiologically relevant molecules and are small in size (5-30 nm). However, their lower electron density makes them poorly visible in electron micrographs, in contrast to the excellent EM visibility of fBSA-Au. Moreover, quantum dots remain invisible in cryosections due to the negative contrasting protocol.

A limitation in the development of bimodal fiducials is that fluorophores can be quenched when they are in close proximity to colloidal heavy metal particles^64–66^. Usually, spacers are incorporated to prevent this quenching, especially when larger sized metal particles are used, but these probes still suffer from a limited fluorescence signal. Thanks to the synthesis strategy of fBSA-Au, where Alexa-labeled BSA proteins are bound to colloidal gold particles, we achieved high fluorescence signal despite using relatively large gold particles. In our case, the bulk of the BSA molecules provides enough space between the gold and the fluorophores to retain sufficient fluorescence. Additionally, commercially available BSA-Alexa555 is labeled with Alexa555 at a 5:1 molar density (5 moles of dye per 1 mole protein). This high labeling density combined with the bulk of BSA should provide a sufficiently large fraction of unquenched fluorophores visible in FM.

One of the challenges with bimodal probes is to ensure colocalization of fluorescence signal and the electron dense particle^44,66^. For fBSA-Au^5^, the challenge primarily lies in the degradation of BSA as it is transported to lysosomes, the enzymatically active end-point of the endocytic pathway. Degradation of BSA will lead to a dissociation of the Alexa label and gold particles, which may lead to labeling discrepancies in correlative approaches. In previous studies, degradation of BSA was seen by the aggregation and clustering of gold particles in lysosomes^34,67^. Similar clustering was observed in lysosomes in the experiments here, indicating the degradation of BSA-Alexa555 (Figures 3 and 4). Our CLEM experiments show no labeling discrepancies between FM and EM; gold particles were seen in every fluorescent-labeled compartment, and vice versa, fluorescence was detected in all compartments containing gold colloids. This overlap confirms that at least within a period of 3 hours uptake the localization and fluorescence of the Alexa dye is retained in endocytic compartments, even after the degradation of BSA, and throughout the process of fixation, sectioning and labeling.

In cryo-EM, accurate selection of ROIs for cryo-ET is of extreme importance due to the fragile nature of frozen hydrated material^68^. The use of fluorescence to determine ROIs prior to imaging is rapidly gaining traction thanks to maturing cryo-FM setups^69–72^. Fluorophores generally retain their fluorescence and exhibit reduced bleaching under cryogenic temperatures^70,73,74^. We have shown that fBSA-Au is compatible with cryo-FM and cryo-EM of vitrified material. By using the 3D distribution of endocytosed fBSA-Au we can select ROIs by cryo-FM, use this fluorescence both to identify where to make lamella in the cryo-FIB-SEM and to locate the targeted organelles within the lamella during subsequent imaging with cryo-ET. fBSA-Au can be a highly suitable fiducial marker especially to use in integrated FM in FIB-SEM systems, where the region of interest (e.g. lamella) identified by FM can be directly prepared for further cryo-ET^75–77^. An additional benefit of endocytosed nanogold fiducials for cryo-ET was also recently reported by providing an accurate tracking during tilt series acquisition and improved tilt-series alignment for the image reconstruction^78^.

Summarizing, we conclude that fBSA-Au is a powerful and easy-to-use 3D fiducial marker that can be used in an array of CLEM applications since it is stable, efficiently endocytosed and compatible with a variety of established LM and EM techniques. fBSA-Au outperforms the existing fiducials with its abundant yet compartmentalized intracellular distribution, and its well-defined small size not obscuring any morphological details. fBSA-Au provides high correlation accuracy both in 2D and 3D CLEM applications, and is especially suited to address questions at the subcellular level requiring nanometer retraction efficiency.

## Materials and methods

### Antibodies and reagents

For immuno-fluorescence we used mouse anti-human LAMP-1 (CD107a) monoclonal antibody from BD Pharmingen (Vianen, The Netherlands), mouse anti-human CD63 monoclonal antibody, clone CLB-gran/12,435 (Sanquin reagents, Amsterdam, The Netherlands) and rabbit anti-human EEA1 antibody (Cell Signalling, C45B10). Primary antibodies were detected with secondary fluorescently (Alexa488/647) labeled antibodies, purchased from Thermo Scientific. Protein A gold 10 nm was made in-house (Cell Microscopy Core, UMC Utrecht, The Netherlands). Alexa555-conjugated BSA was bought from Thermo Scientific (A34786). Tannic acid was purchased from Mallinckrodt. Chloroauric acid trihydrate was purchased from Merck (#1.01582). Paraformaldehyde (95% wt/vol) was purchased from Sigma. Glutaraldehyde (8% wt/vol; EM grade) was purchased from Polysciences.

### fBSA-Au^5^ and fBSA-Au^10^ complex synthesis

5 or 10 nm colloidal gold particles were synthesized by reduction of chloroauric acid with tannic acid and sodium citrate, according to protocols developed by Slot and Geuze ^31^. Reagent ratios were adjusted to obtain 5 or 10 nm sized colloid gold particles. Following synthesis, the colloid particles were stabilized with an excess of AlexaFluor 555-labeled BSA, as described previously for other proteins^30,31^. For additional stabilization, 0.1% BSA (final concentration) was added to the solution. The complexes were centrifuged on a 10-30% glycerol gradient centrifugation to remove aggregates and excess protein. The purified fraction was diluted in PBS and stored with the addition of sodium azide. Prior to use in cell culture, the required volume of fBSA-Au was dialyzed against PBS overnight at 4°C to remove the sodium azide and residual contaminants.

Measurements for sizing of synthesized gold colloids were performed by diluting gold colloids or fBSA-Au at OD5 in dH_2_O. Formvar and carbon-coated copper grids were placed on 5 μl drops of diluted solutions for 5 minutes. Grids were washed once on drops of dH_2_O, after which excess liquid was drained using filter paper. After drying, the gold particles were then imaged in TEM at magnifications >80,000×. Micrographs containing gold particles were analyzed using the particle size analyzer developed by Ralph Sperling^79^. At least 100 gold particles are measured per condition except for non-functionalized Au^10^ (n=46). Data is represented as mean ± S.D (Figure 1).

### Cell culture

Hela and U2OS cells were cultured in Dulbecco’s Modified Eagle’s Medium (DMEM; Gibco) supplemented with 10% heat-inactivated fetal bovine serum (FBS), 2mM L-glutamine, 100 U/ml penicillin, 100 μg/ml streptomycin (complete DMEM). Cells were grown under 5% CO_2_/air atmosphere at 37°C.

Primary dorsal root ganglion (DRG) neuron cultures were derived from spines of 6-8 week old wild-type mice after CO2 inhalation and exsanguination. Experiments were licensed under the UK Animals (Scientific Procedures) Act of 1986 following local ethical approval. All procedures were carried out in accordance with UK Home Office regulations. DRG from each spine were isolated and kept at 4°C in HBSS (Thermo Fisher) supplemented with 20 mM Hepes pH 7.4 (‘HBSS+H’). The ganglia were washed twice by pelleting at 800g for 3 min and resuspending in 5 mL ‘HBSS+H’ then enzymatically digested by resuspension in 1 mL 37°C HBSS supplemented with 15 μL 20 mg/mL collagenase type IV (Thermo Fisher). After 1 hour incubation at 37°C, 5% CO_2_, 1 mL pre-warmed HBSS supplemented with 100 μL 2.5% Trypsin (Gibco) was added. After 15 minutes, 5 mL ‘plating media’ containing Neurobasal media (Thermo Fisher), 1 x B-27 (Thermo Fisher), 2 mM L-Glutamine (Thermo Fisher), 5% FBS (Thermo Fisher), 20 mM Hepes pH 7.4, 100 U/mL penicillin and 100 U/mL streptomycin was added. Trituration was performed using a 1mL pipette after washing twice in 2 mL ‘plating media’. The resulting cell suspension was layered onto a 4°C, 3 mL 15% BSA cushion (BSA in DMEM) and spun at 4°C, 300g for 8 min. The cell pellet was resuspended in 0.5 mL pre-warmed ‘plating’ media supplemented with 100 ng/μL NGF (Peprotech). Cells from all spines were pooled before plating in microfluidic devices (MFDs) or on cryo-EM grids. Next day, media was replaced with maintenance media containing Neurobasal media, 1 x B-27, 2 mM L-Glutamine, 20 mM Hepes pH 7.4, 100 U/mL penicillin, 100 U/mL streptomycin, 100 ng/μL NGF and 40 μM 5-UfDU (uridine and 5-fluorodeoxyuridine, Sigma). Cultures were maintained at 37°C, 5% CO_2_ and half media replaced every 2-3 days.

### Immunofluorescence labelling and imaging of endocytosed fBSA-Au

Cells grown on glass coverslips were treated with fBSA-Au^5^ in culture medium at OD5, and incubated for 3 hours at 37°C. Cells were fixed with 4% formaldehyde in PBS for 1 hour, and permeabilized with 0.1% Triton X-100 in PBS. Blocking was performed using 1% bovine serum albumin (BSA) in PBS. Immunolabeling for LAMP-1 and EEA-1 was performed by incubating coverslips in PBS containing the corresponding antibodies and 1% BSA. Labeling was visualized using Alexa-tagged secondary antibodies. After secondary labeling, coverslips were washed with PBS and dH2O, and mounted to microscope slides using Prolong Gold or Diamond (Thermo Scientific).

### Correlative microscopy of resin-embedded samples

For correlation of fluorescence microscopy and EM of resin-embedded cells, imaging was performed prior to sample preparation in EM. Cells were grown on carbon-coated, gridded coverslips prepared as in ^46^, and treated with fBSA-Au^5^ diluted to OD5 in complete DMEM for 3 hours. Cells were washed in 1× PHEM buffer to remove excess fBSA-Au^5^, and fixed using 4% formaldehyde and 0.2% glutaraldehyde in 1× PHEM buffer. Using FM, Z-stacks of cells of interest were obtained for the Alexa555 signal. The position of cells relative to the pattern etched in the coverslip was registered using polarized light.

To prepare specimens for electron microscopy, the imaged coverslips were postfixed using osmium tetroxide and uranyl acetate, dehydrated using a graded ethanol series, and embedded in Epon resin. Resin was polymerized for 48 hours at 65°C. After polymerization, the glass coverslip was removed from the Epon block by dissolving it in hydrogen fluoride, after which the exposed Epon surface was thoroughly cleaned with distilled water and left to harden overnight at 63°C. Areas of the resin block containing imaged cells were cut out using a clean razor blade, and glued to empty Epon sample stubs, with the basal side of the cells facing outwards. From these blocks, 70 and 250 nm thick sections were cut and collected on formvar and carbon coated copper support grids (50 mesh or slot grids). Grids with 250 nm thick sections were seeded with tomography fiducials by placing the grids on drops of ddH2O containing 1:100 diluted protein-A-gold 10 nm for 5 minutes. Afterwards, grids were rinsed 3 times on distilled water and blotted dry with filter paper.

### Sample preparation and light microscopic imaging of Tokuyasu cryosections

For CLEM on thin (70 nm) and thick (350 nm) cryosections, cells were grown in 60 mm culture dishes, treated with fBSA-Au^5^ diluted to OD5 in complete DMEM for 3 hours at 37°C and fixed with 2% formaldehyde and 0.2% glutaraldehyde in 0.1M phosphate buffer (pH 7.4). Samples were gelatin embedded, cryoprotected, sectioned and immunolabeled according to previous protocols^2,9^, with minor modifications. Following incubation with primary antibodies, the grids were labeled with Alexa488 labeled secondary antibodies, followed by incubation with protein-A gold conjugates (10 nm). The grids were washed with dH2O and placed between a microscope slide and a #1 coverslip in 2% methylcellulose in dH_2_O. Sections were imaged in a Deltavision RT widefield FM (GE Healthcare, U.S.A.) equipped with a Cascade II EM-CCD camera (Photometrics, U.S.A.). Grids were first imaged at 40× magnification to form a map of the section, after which regions of interest were selected using 100× magnification. After imaging the grids were removed from the microscope slide, thoroughly rinsed with H_2_O and contrasted for EM and embedded in methylcellulose containing uranyl acetate, according to previous protocol ^2^.

### Electron microscopy of resin sections and Tokuyasu cryosections

Thin cryosections were imaged in a Tecnai 12 TEM (Thermo Fischer Scientific, Eindhoven, The Netherlands) equipped with a Veleta 2k×2k CCD camera (EMSIS, Munster, Germany), operating at 80 kV. Tilt series of resin sections and labeled thick cryosections were acquired in a Tecnai 20 TEM (Thermo Fischer Scientific) operating at 200 kV, equipped with an Eagle 4K×4K CCD camera running Xplore3D (Thermo Fischer Scientific) software. Single tilt image series were automatically collected with 1° tilt increments from −60° to +60° at microscope magnifications of 11500× or 14500×, resulting in final pixel sizes of 0.96 nm or 0.76 nm, respectively.

### Cryo-FM and Cryo-EM of DRG Neurons

For preparation of EM grids, a thin layer of homemade continuous carbon was floated on top of individual Quantifoil R3.5/1 200 mesh gold grids (Quantifoil Micro Tools). After drying, these were plasma cleaned using Nano Clean Plasma cleaner Model 1070 (Fischione) for 40 s at 70% power in a 9:1 mixture of Argon and Oxygen gas. Grids were transferred into a Ibidi μ-slide 2 well co-culture dish (Ibidi) and coated with poly-L-lysine and laminin as described for the MFDs. After growth of DRG for 4 DIV, fBSA-Au^5^ was prepared as for live imaging and incubated for 2 hours. Samples were washed twice in pre-warmed maintenance media lacking 5-UfDU then vitrified by plunge freezing into liquid ethane after manual back-side blotting in a Vitrobot Mk II (Thermo Fisher) kept at 37°C, 100% humidity.

Cryo fluorescence microscopy was performed using a Leica EM cryo-CLEM wide-field microscope (Leica Micosystems) equipped with a 50x/0.90 NA DRY cryo-objective lens. 15 μm z-stacks of the entire grid with 1 μm z-spacing were acquired using the Leica LAS X Matrix software in green, red and transmitted light channels. Correlation was performed manually during cryo-EM imaging. Cryo-electron tomograms of the thin parts of DRG axons were acquired using a TITAN Krios G3 (Thermo Fisher) operated at 300kV equipped with K3 detector and Quantum GIF (Gatan) with slit width 20 eV. Tilt series were acquired using SerialEM^80^ from ±60° in 2° increments using a dose-symmetric scheme^81^ with defocus set from 3.5 - 6 μm underfocus. The dose in each image was 2e^-^/Å^2^ with pixel size 2.68Å/pix, leading to total dose 120e^-^/Å^2^. Movies were acquired in counting mode, with 10 frames per tilt image.

### Cryo-FM and Cryo-EM of U2OS cells

Gold grids (Quantifoil, R2/2) were glow-discharged and incubated on top of 40 μL droplets of fibronectin (50 μg/mL) for 2-3 hours at 37 °C. 90.000 U2OS cells were seeded on these grids in 30 cm glass bottom dishes (Greiner bio-one). After 72 hours, 3 μL of 1 μm Dynabeads (Thermo Fischer Scientific: MyOne with 40% iron oxide, carboxylic acid) diluted 1:10 in PBS, was added to the grids and the cells were vitrified in liquid ethane after manually blotting for 12s. Prior to plunging, the cells were incubated at 37°C with fBSA-Au^5^ diluted to OD 5 forOD5 for 3 to 4 hours.

fBSA-Au^5^ spots particleswere localized in the FEI CorrSight™, a spinning-disk confocal light microscope equipped with a cryo-module. Grid overview images were taken with a 5x/0.16 NA air objective. 10 μm z-stacks of individual cells with 300 nm spacing were obtained with a 40x/0.9 NA air objective after excitation at 488 (Dynabeads) and 561 (fBSA-Au^5^) nm. Image acquisition was done using FEI MAPS 3.8 software. Next, the grid was loaded in the cryo-FIB (Aquilos™, Thermo Fisher Scientific), the SEM grid overview image was correlated to the light microscope overview image in MAPS3.8 using the 3-point alignment method. Individual z-stacks were correlated to the SEM image (1536×1024, 1 μs, 2 kV, 13 pA) of target cells, prior and post milling. The Dynabeads were used as fiducials to perform the x-y-z correlation between the SEM images and the z-stacks. Transformation parameters were determined using the 3D Correlation Toolbox and the transform was applied using Pyto, a python-based package for cryo-ET analysis, as described in Arnold *et al*.^48^.

Lamellae were prepared as described in Wagner *et al*.^54^, at 16 degree stage tilt with a stepwise decreasing current of 1 to 0.3 to 0.1 nA. The final polishing step was performed at 30 pA to reach a lamella thickness of 100-200 nm. Lamellae were imaged on a 200 kV Talos Arctica transmission electron microscope (Thermo Fisher Scientific) with a post-column energy filter (slit width of 20 eV) and a K2 summit direct electron detector (Gatan). Lamellae overview images were recorded at a pixel size of 18.72 Å/px. The lamellae visible in the previously recorded post-milling SEM images were overlayed in FIJI with the lamella overview images recorded in the cryo-TEM to allow correlation of the LM data to the cryo-TEM lamella overview images and thus the localization of the beads in the lamellae. High magnification tilt series of the beads were recorded using SerialEM at a pixel size of 2.17 Å/px, a dose rate of ^~^3 e^-^/px/s and a total dose of 100.3 e^-^/Å^2^. The tilt series were collected with a 2° tilt increment from +69° to −51° at a defocus of 3.

### Tomogram reconstruction

Tomogram reconstruction was performed using the IMOD software package^41^. Tilt series from ET were aligned using bead tracking of the 10 nm gold particles used for immunolabeling (cryosections) or seeded gold fiducials (resin sections). Tomograms were generated from the aligned data using weighted back projection. For DRG neurons, gain correction, CTF-estimation and motion correction were performed using the IMOD programmes alignframes and ctfplotter. Fiducial model generation, tomogram alignment and reconstruction was performed in eTomo. The resulting volumes were binned by 4 and filtered for visualization using the deconvolution filter implemented in WARP. The tilt series from U2OS cells were aligned and dose-weighted using MotionCor2 ^82^. Alignment of the tilt series via either fiducial or patch tracking, and tomogram reconstructions were performed in eTomo, part of the IMOD/4.10.29 package ^41^. After CTF correction was performed in IMOD, the tomograms were reconstructed using weighted back projection and a SIRT-like filter. Tomograms were 4x-binned and low pass filtered to 40 Å for visualization using the TOM Toolbox ^83^.

### Correlation of light and electron microscopy images

Registration of thin section fluorescence and EM data was performed using ec-CLEM^40^. Here, multiple corresponding pairs of fluorescent spots and gold particles were manually selected, after which the software automatically applies the correct scaling and transformation steps and generates overlays of FM and EM data. We used only linear transformation options to achieve the overlays shown in the Figures.

For correlation of FM and ET data, registration was first performed using ec-CLEM by overlaying FM data over a regular TEM image of the ROI, collected before the start of tilt imaging. The transformed fluorescence images were then overlaid with the tomogram slices corresponding to the region of the regular TEM image.

## Supporting information

Supplementary Video 1: Live-cell imaging in HeLa cells

Supplementary Video 2: Tomogram of Figure 6C

Supplementary Video 3: Tomogram of Figure 6F

Supplementary Video 4: Tomogram of Figure 6G

Supplementary Video 5: Cryo-FM Z-stack of Figure 7

Supplementary Video 6: Tomogram of red arrow in Figure 7

Supplementary Video 7: Tomogram of a microtubule-late-endosome connection

## Acknowledgments

This work was supported by funding from Stichting voor de Technische Wetenschappen (STW), grant number 12715, granted to HG and JK. NL also acknowledges the Netherlands Organization for Scientific Research (NWO) for an ZonMW-TOP grant awarded to JK [40-00812-98-16006]. The electron microscopy performed within this work is partially subsidized by the research programme National Roadmap for Large-Scale Research Infrastructure (NEMI) of NWO [184.034.014]. Cell Microscopy Core, UMCU is part of the Dutch Correlative Light Electron Microscopy node of EuroBioImaing.

## Author Contributions

NL, JF, GP, HG, and JK designed the study. GP conceived the bimodal probes. TV, VO, SvanD synthesized, optimized and characterized the endocytic fiducials. JF, TV, LY and CdeH optimized the cellular uptake and imaging of endocytic fiducials and analyzed the data with NL. JF performed 2D on-section CLEM and 3D correlative FM – ET studies. HF performed cryo-ET experiments of neurons. LdeJ and SH performed cryo-CLEM experiments of U2OS cells. JF, LdeJ, and HF did the image correlation and segmentation, and prepared the corresponding figures with NL. WM, WL, SH, AC, FF, HG, JK and NL supervised the study. JF and NL wrote the paper with input from all authors. All authors reviewed the manuscript.

## Supplementary data

Supplementary Figure 1: Accuracy of registration using fBSA-Au fiducials

**Supplementary Figure 1:**
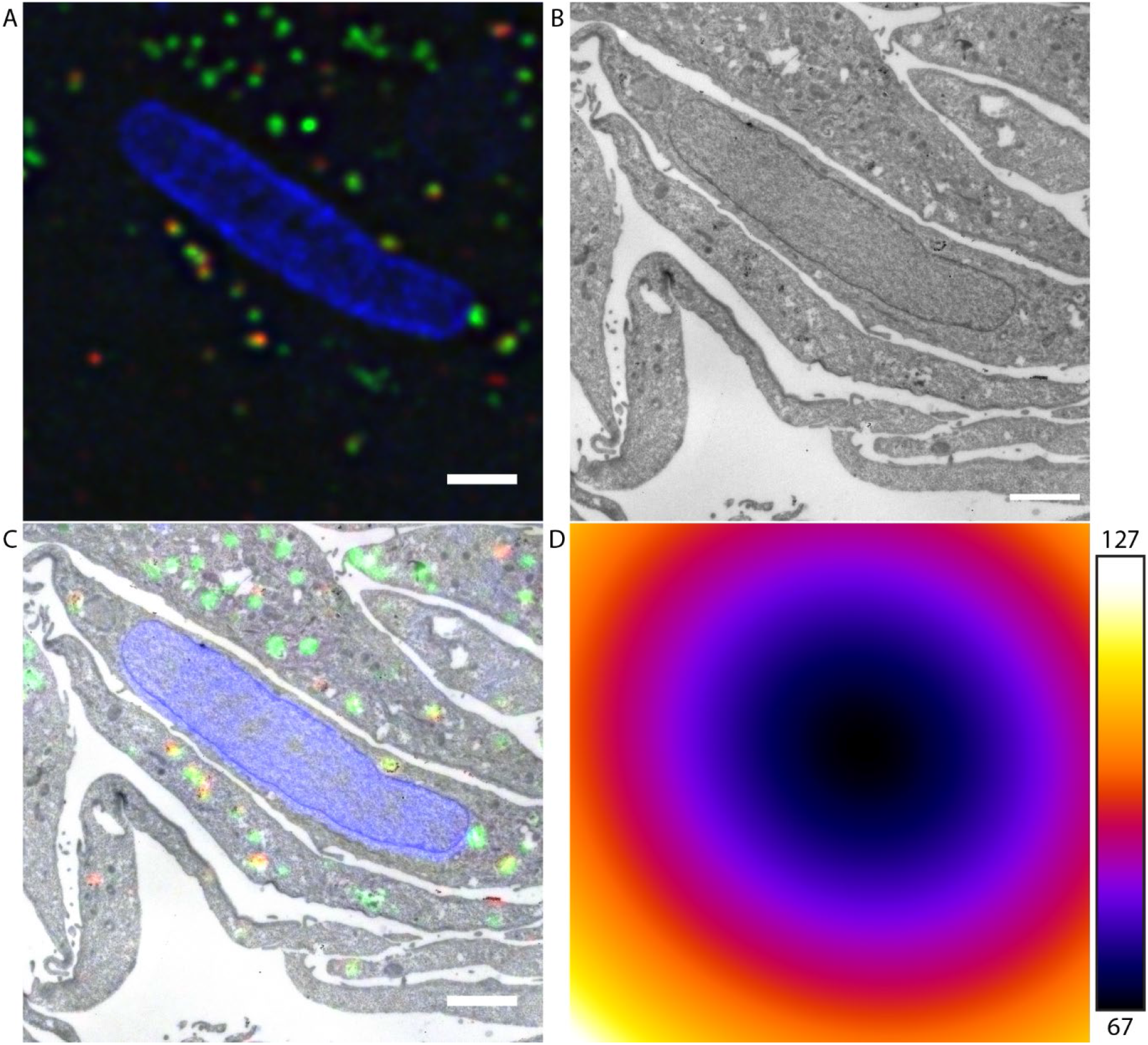
fBSA-Au as fiducial enables high registration accuracy. (A) FM data of region of interest. (B) EM region of interest. (C) Overlay of FM and EM data. (D) Quantification of registration error using ec-CLEM, showing regions of high accuracy (67 nm registration error) and lower accuracy (127 nm registration error). *Scalebars: 2 μm*.

## List of Supplementary Videos

1. Supplementary Video 1: Live-cell imaging in HeLa cells
2. Supplementary Video 2: Tomogram of Figure 6C
3. Supplementary Video 3: Tomogram of Figure 6F
4. Supplementary Video 4: Tomogram of Figure 6G
5. Supplementary Video 5: Cryo-FM Z-stack of Figure 7
6. Supplementary Video 6: Tomogram of red arrow in Figure 7
7. Supplementary Video 7: Tomogram of a microtubule-late-endosome connection

